# Facile assembly of an affordable miniature multicolor fluorescence microscope made of 3D-printed parts enables detection of single cells

**DOI:** 10.1101/592170

**Authors:** Samuel B. Tristan-Landin, Alan M. Gonzalez-Suarez, Rocio J. Jimenez-Valdes, Jose L. Garcia-Cordero

**Affiliations:** Unidad Monterrey, Centro de Investigación y de Estudios Avanzados del IPN, Parque PIIT, Apodaca, Nuevo León, CP. 66628, Mexico

## Abstract

Fluorescence microscopy is one of the workhorses of biomedical research and laboratory diagnosis; however, their cost, size, maintenance, and fragility has prevented their adoption in developing countries or low-resource settings. Although significant advances have decreased their size, cost and accessibility, their designs and assembly remain rather complex. Here, inspired on the simple mechanism from a nut and a bolt we report the construction of a portable fluorescence microscope that operates in bright field mode and in three fluorescence channels: UV, green, and red. It is assembled in under 10 min from only six 3D printed parts and basic electronic components that can be readily purchased in most locations or online for US $85. Adapting a microcomputer and a touch LCD screen, the microscope can capture time-lapse images and videos. We characterized its resolution and illumination conditions and benchmarked its performance against a high-end fluorescence microscope by tracking a biological process in single cells. We also demonstrate its application to image cells inside a microfluidic device in bright-field and fluorescence mode. Our microscope fits in a CO_2_ chamber and can be operated in time-lapse mode. Our portable microscope is ideal in applications where space is at a premium, such as lab-on-a-chips or space missions, and can find applications in clinical research, diagnostics, telemedicine and in educational settings.

## Introduction

Fluorescence microscopy is an essential tool in biomedical research used to visualize, analyze and study molecules, cells and tissues. One of its main applications is to enable the quantification and localization of the distribution of cellular components [1], which enables quantitative biology. Other important applications include its usage as a readout mechanism in biochemical assays such immunoassays or qPCR. In the healthcare sector, fluorescence microscopy has been recommended by the World Health Organization for the diagnosis of tuberculosis [2]. However, because of its cost, training, maintenance and fragility, conventional fluorescence microscopes remain out of reach in developing countries, in rural areas and in remote settings [3–6]. Thus, access to affordable fluorescence microscopes (albeit at the cost of compromising certain functionalities and resolution) can facilitate its deployment in these settings and its usage in point-of-care diagnostics, telemedicine, and environmental monitoring, benefitting global healthcare. Furthermore, producing affordable and easy-to-assemble laboratory tools and instrumentation could facilitate research, produce quality results [7], improve laboratory productivity [8], and can potentially lead to new discoveries in biology, physiology, chemistry, and biomedicine [9,10].

Affordability and portability in fluorescence microscopy has been accomplished by retrofitting optical elements (objectives, filters, LEDS, lasers, lenses, mirrors) and 3D-printed parts to smartphones cameras [3,5,11–13]. However, smartphones, although extremely powerful and unique, are difficult to reconfigure and take apart [14]; evolve at a rapid pace which may render them obsolete; constant software updates on the operating systems may disrupt their performance [15]; their size hampers further miniaturization and interferes in the integration with other additaments or sensors; and are not optimized for long-term biological experimentation [16]. A different approach has been to use regular cameras (webcams or digital cameras) placed in a framework made of plastic parts, metallic hardware, and 3D-printed parts [5,6,16,17]; however, in some cases, the complexity of their designs encumbers downstream integration with other 3D printed parts and the amount of pieces needed to put it together may be overwhelming for a newcomer or for technologists trying to develop affordable instrumentation in low-resource settings. It also hampers its prompt adoption by scientists who are setting up a laboratory with limited resources and who need a multi-channel fluorescence microscope with basic functionalities and sufficient quality for most biological applications.

To solve some of these issues, we introduce a low-cost and portable multicolor fluorescence microscope the size of a cube with a side length of 3 cm. Its operation and framework are inspired in the mechanism of a nut and bolt. The microscope is assembled from only six 3D-printed parts, 4 power LEDs, an 8MP CMOS camera, a metallic rod, and a PCB with basic electronics. Aside from the Foldscope [18] (which does not contain any camera sensor), our microscope is assembled using the least amount of pieces possible. After pieces are 3D-printed and the PCB manufactured, the microscope can be assembled in 10 min (see **S1 Video**) with the capability to acquire images in brightfield and 3 color fluorescence channels. We present its operation, characterization of its illumination homogeneity in each of the fluorescence channels, and its demonstration in a cellular biological assay where it is possible to track single cells captured in microwells.

## Materials and methods

### 3D-Printing Chassis

Microscope parts were designed in Inventor (ver 2017, Autodesk) and fabricated in a 3D printer (Makerbot Replicator 2); **see S1 Fig**. Dimensions of the microscope tube are shown in **S2 Fig**. The parts are made of black polylactic acid (PLA) and printed with the following settings: speed in X and Y axis of 40 mm/s, temperature of 230°C, layer thickness of 200 μm, and infill of 10%. Once fabricated, pieces were manually assembled; see **S1 Video** for a step-by-step assembly.

### Electronics and optical components

The optical system relies on the Raspberry Pi Camera Module V2 that comes with an 8-MegaPixel CMOS image sensor camera and a plastic lens. This lens comes with the camera and no further specifications are given on it; however, US Patent 7,564,635 B1 provides the description of a lens that has the same focal length (3.04 mm) provided in the specification of this camera module. High brightness LEDs (white, UV, green and blue) were used to illuminate the sample. Plastic color filters (Roscolux) or single-band pass optical filters (Zeiss) were used as emission filters and placed between the image sensor and the lens. A custom printed circuit board (PCB) containing the electronic components to control the LEDs were placed below the camera. The schematic and the list of components can be found in **S3 Fig** and **S1 Table**, respectively.

### Graphical User Interface (GUI)

A single-board computer (Raspberry Pi) was used to control the LEDs and to program a graphical interface using Python and OpenCV. The GUI allowed to acquire images and control different parameters such as the exposure time (**S4 Fig**).

### Cell assays

To compare image quality acquired with our microscope and a conventional microscope, THP-1 cells were permeabilized with Tween-20 (P1379, Sigma Aldrich) 0.05% in PBS 1X for 15 min, centrifuged at 1,200 rpm for 5 min, and washed with PBS 1X. Next, cells were either incubated for 20 min either with DAPI (2.9 μM, 62247, Thermo Fisher Scientific) or Calcein-AM (20 μM, C1359, Sigma Aldrich), or for 10 min with Ethidium Homodimer (16 μM, EthD-1, E1903 Sigma Aldrich). The cells were washed again with PBS 1X and centrifuged at 1200 rpm for 5 min. Then, 20 μL of the cellular suspensions were spread over three different clean coverslips, allowed to dry, and incubated with 10 μL of glycerol. Finally, a second coverslip was placed on the top of the samples. For Calcein-AM stained cells, PBS 1X was used instead of glycerol. Finally, both coverslips were sealed with enamel.

### Neutrophil assay

Human peripheral-blood neutrophils were purified from human blood by a non-continuous Percoll density gradient. After purification, neutrophils nuclei were stained with Hoechst 33342 (H3570, Thermo Fisher Scientific) at 16.2 μM for 5 min. Neutrophils were stimulated with either Hank’s solution (control) or with *E. coli* lipopolysaccharide (LPS at 100 μg/mL, L2755, Sigma Aldrich). Both solutions contained also Sytox Orange (S11368, Thermo Fisher Scientific) at 5 μM for nucleic acid staining. See Supplementary Information for further details. Cells were placed in three PDMS-devices containing microwells (20 μm diameter, 20 μm height). One of the devices was placed on the miniature microscope with the stimulus solution and the other two on an inverted fluorescence microscope (Axio Observer, Zeiss). 20 μL of the cell suspension was added to the three devices. After 5 min, 20 μL of the stimulus were added to two devices, one in our microscope and the other in the Zeiss microscope. 20 μL of Hank’s solution was added to the other device sitting on the Zeiss microscope, which served as a negative control. Next, images in brightfield and red and blue fluorescence channels were acquired every 10 min in both microscopes. Images were analyzed using an image processing software (Fiji) where fluorescence intensity was measured over time for all individual microwells.

### Cell tracking in a microfluidic device

A one-channel microfluidic device was designed in AutoCAD (Student version, Autodesk) and fabricated by soft-lithography [19]. The device consists of an inlet, an outlet, and a long serpentine channel with a height and width of 20 and 40 μm, respectively. The device was placed on the miniature microscope stage and a THP-1 cell suspension, stained with Calcein-AM, was flowed into the channel using a 1 mL syringe. The center of the device was focused, and a video was recorded in bright-field and in the green fluorescence channel while the cells were flowing through the serpentine channel.

### THP-1 culture and time lapse microscopy

THP-1 cells were incubated using supplemented RPMI-1640 medium (11875093, Thermo Fisher Scientific) inside a T25 flask before experimentation. A 1 mL chamber was fabricated using a PDMS slab, plasma bonded to a cover slide and exposed to UV light for 3 hours. 900 μL of fresh media was added to the chamber followed by 100 μL of THP-1 cell suspension at 100×10^5^ cell/mL. Another block of PDMS was used to seal the chamber from the top. The cell chamber was placed on the miniature microscope stage and the whole microscope inside a cell culture incubator at 37°C with 5% CO_2_. The bottom of the chamber was manually focused, and the miniature microscope was set in time-lapse mode on the GUI. Next, all cables were disconnected, except for the power cord. During the first ∼24 hours, brightfield images were acquired every 15 min. For the last ∼2 hours, images acquisition was set to every minute.

## Results and discussion

### Microscope Design

Our goal was to engineer a miniature multicolor fluorescence microscope of simple design that could be built from the least number of pieces and thus be assembled rapidly, and which, while using 3D printed pieces, could still offer enough accuracy to focus on objects while being mechanically stable and indifferent to vibrations. Importantly, our goal was not to develop a fluorescence microscope that offered the same illumination and image quality as a commercial microscope but rather a microscope that could be assembled in a short period of time, was affordable for low-budgets labs, offered sufficient image quality for most applications (with enough resolution to image single cells) and enable quantitative biological measurements, and equally important, that it was simple enough for any scientist to observe biological objects in brightfield and stained with multiple fluorescent dyes.

The microscope is inspired in the mechanism of a nut and bolt, **Fig 1a**. The bolt head serves as a base and as a casing to enclose the sensor camera, while the tip of the bolt holds the plastic lens, **Fig 1b,c**. The nut functions as a stage that fits on the threaded shank, supports the samples, and serves as a lid for fluorescence observations (**Fig 1d**). The threading in the shank is 1 mm (**S2 Fig**). It is important to realize that because the microscope is made of PLA, the threading can wear out over time and thus decrease its ability to finely focus. The microscope base (bolt) is fixed while the stage rotates to bring the object into focus. One disadvantage of this arrangement is that the object under observation is rotated with the stage, yet once focused the object can be manually lifted from the stage and rotated to preserve the orientation.

**Fig 1.**
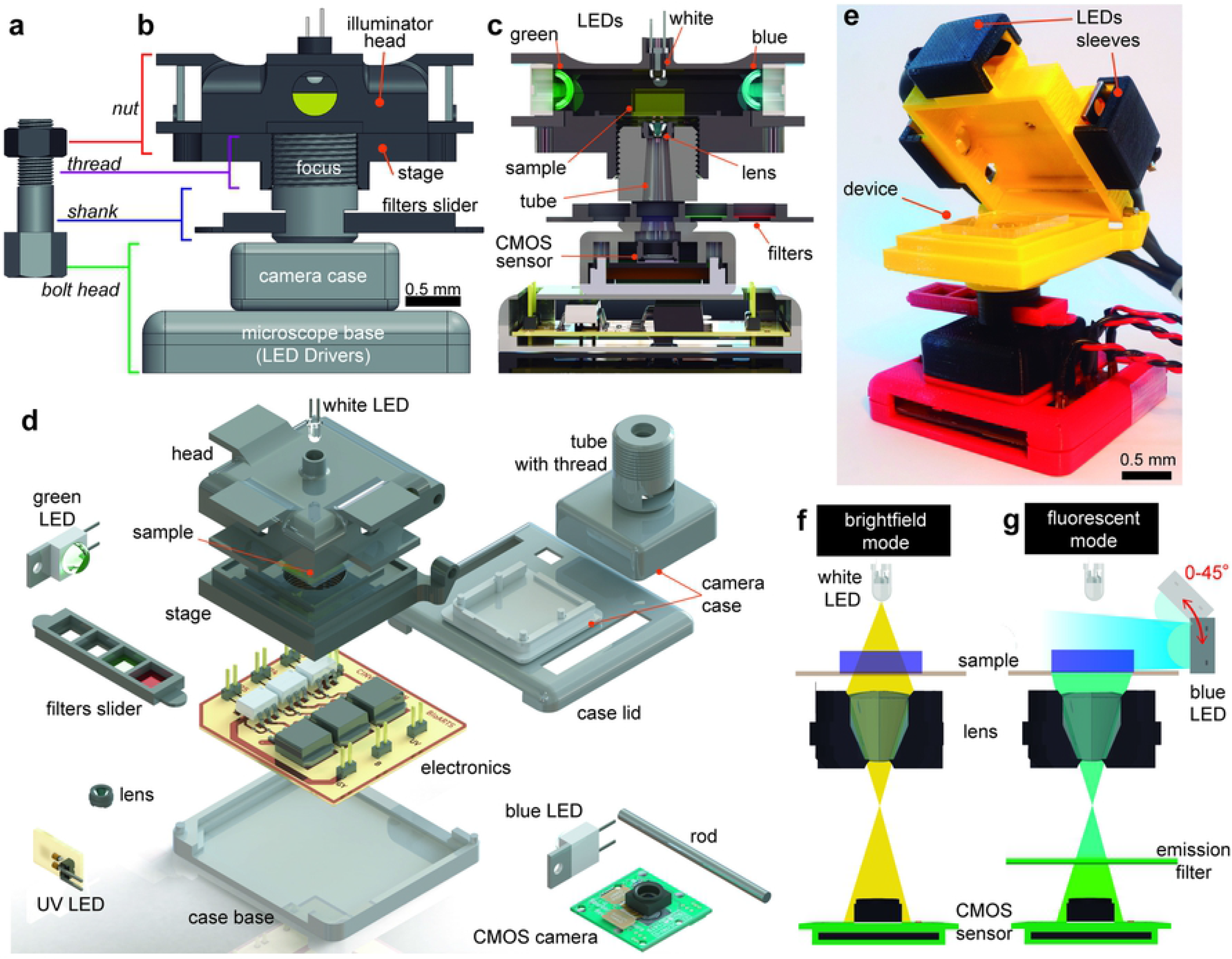
Anatomy of the miniature microscope. (**a**) The mechanism of the microscope is inspired by how a nut and bolt works. (**b**) Front view of the microscope. (**c**) Cutaway diagram of the microscope. (**d**) Exploded view of the different components comprising the microscope. (**e**) Photograph of the microscope with the lid open and a microfluidic device mounted on the stage. Schematic of the light path during operation of the microscope in (**f**) brightfield and (**g**) fluorescence mode. Color LEDs can be placed at different angles (0-45°) to provide a homogeneous illumination.

Most fluorescence microscope configurations make use of fluorescence detection cubes (exciter, dichroic, and emitter) incorporated in a filter wheel. Because LEDs have a very narrow band spectrum (20-40 nm) it is possible to omit the detection cube and just keep the emission filter. Thus, the stage performs as a lid that enables brightfield observation using a white LED placed on the center top of the lid (**Fig 1e**) and allows fluorescence measurements using 3 high-power color LEDs (UV: 385 nm, Green: 475 nm, and Blue:535 nm) that illuminates the sample from the sides (**Fig 1f**) Three plastic emission filters are placed on a hollowed-out tray that slides on the shank of the microscope and in which one of the hollows remains empty for brightfield illumination. With a small modification to the shank, regular microscope optical filters can be also employed. The simplicity of our configuration contrasts with other microscopes that require several 3D-printed parts [17] or acrylic parts, nut, bolts, and tubes, for their assembly [16,20].

### Microscope Parts

The microscope objective consists of a plastic lens detached from a commercial camera module and placed upside-down 25.4 mm above an 8 MP CMOS sensor (**Fig 1e**), in a similar configuration to what others have reported [16,21,22]. The sensor pixels contain three color pigment mosaic filters (Red, Green, Blue) that serve as extra emission filters by digitally extracting each channel from the raw image [16].

Our microscope is assembled from six 3D-printed pieces (a lid, a stage, a shank, front and back casing, and a tray), a metallic rod, 4 high-power LEDs, a CMOS camera, a lens, and an electronic control unit that powers the LEDS from a single 9V power supply, see **S1 and S3 Figs**. The 3D printed pieces are fabricated in 2.5 hours while the microscope is assembled under 10 min. The manual step-by-step assembly of the microscope are shown in **S1 Video**. Altogether, the size of the microscope is similar to a cube with a side length of ∼3 cm and weighs only 58 g. The total cost of the opto-electro-mechanical system is $85 (USD), see **S1 Table**. In comparison, a low-entry commercial microscopes with a single fluorescence module can cost up to ∼$1900 and weight ∼9.5 kg [4]. Because of its popularity and low-cost, we selected the microcomputer Raspberry Pi to control the microscope and process the images. An added benefit of the Raspberry Pi platform is the plethora of plug-in sensors and accessories [14] that in the future could improve the capabilities of our microscope and aid in the automation of biological and biochemical assays. Adding the cost of the Raspberry Pi and the tablet, the total cost of our microscope increases to US$202.

### Microscope Operation

To operate the microscope, we created a graphical user interface (GUI) in Python that allowed us to turn on and off on-demand the different color LEDs; set the exposure time; capture still images, videos, or time-lapse images; save images in different formats; among other functions, **S4 Fig**.

**Fig 2** shows the step-by-step operation of the microscope. It involves placing the sample on the stage by opening the lid. Next, the lid is closed to isolate the sample from external light sources. For brightfield observations, the white LED is turned on from the GUI and the tray is shifted to match the empty hole of the tray. The stage is manually rotated clockwise or counterclockwise to focus the objects on the sample. For fluorescence observations, one of the color LEDs are turned on while the filter tray is manually slid to match the corresponding emission filter with its corresponding LED. **S2 Video** shows how easily the microscope is operated by a user.

**Fig 2.**
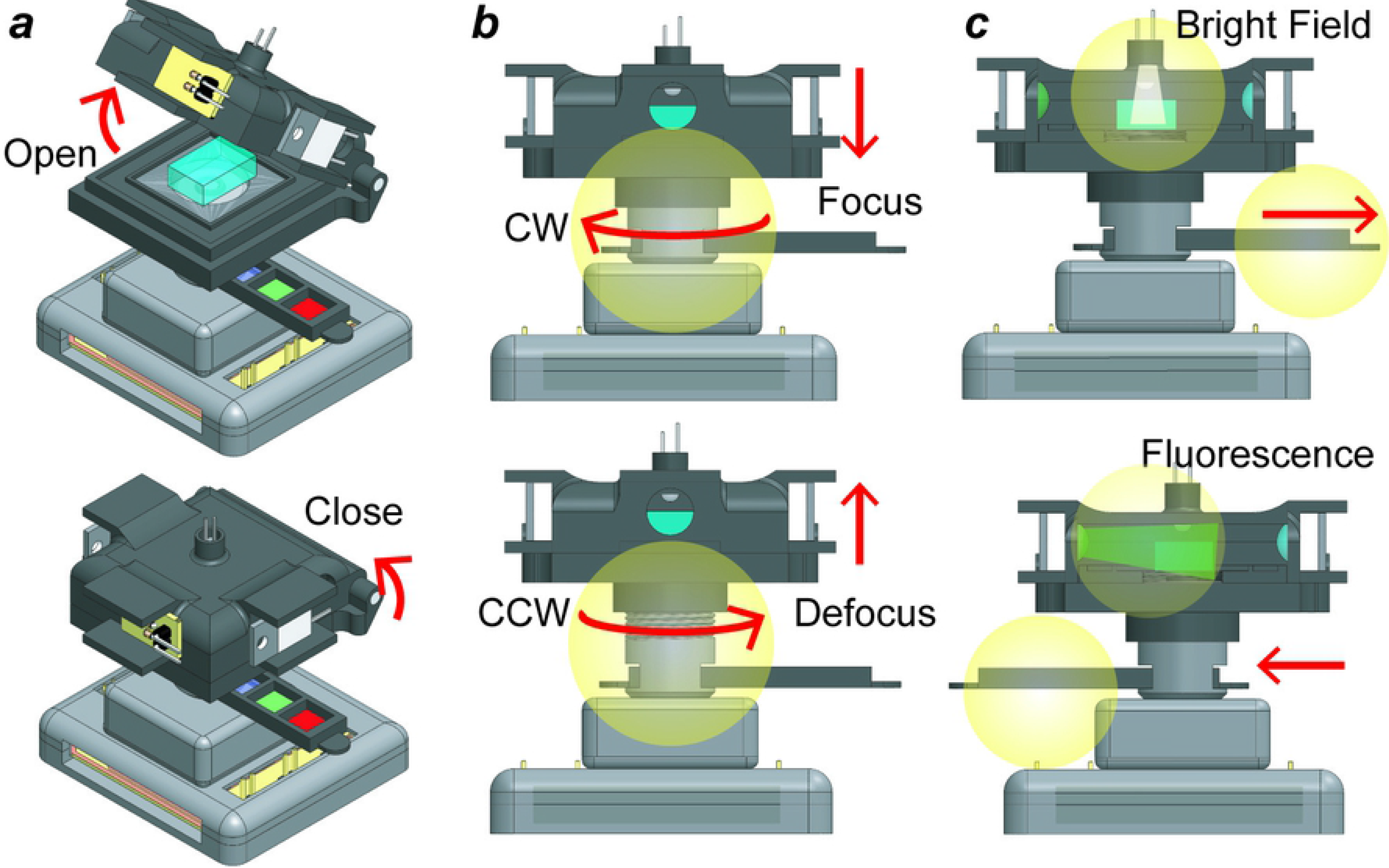
Step-by-step operation of the miniature microscope. (**a**) The lid is open to place a sample and then closed to prevent any external light sources to enter the stage and thus affect the image capture process. (**b**) Rotating the stage clockwise brings the sample closer to the lens, while rotating it counterclockwise moves the sample away from the lens. (**c**) To acquire bright field or fluorescence images, the slider is moved manually either left or right to match the emission filter with the excitation LED. The leftmost hollow in the filter tray is kept empty for bright-field observation.

### Microscope Resolution

The field of view of our setup is ∼0.2 mm^2^ (525 μm × 394 μm) with an estimated equivalent magnification of a 20x microscope objective. To achieve different magnifications the distance from the lens to the sensor can be adjusted. The theoretical optical resolution, calculated at 550 nm according to the Rayleigh criterion, is 0.448 μm. The focal length and working distance of the lens are approximately 2.7 and 0.1 mm, respectively, and the lens has a numerical aperture of 0.45, while the depth of field is 1.46 μm. To determine the optical resolution of our system, we determined the full width at half maximum (FWHM) of the point spread function (PSF) for 40 1-μm fluorescent beads (Firefli Fluorescent Blue) spread over the surface. The estimated FWHM of the 40 microbeads with our microscope is 1.187 μm, identical the one obtained with a 20X objective using a Zeiss microscope, **Fig 3a,b**. We also imaged a 1951-USAF resolution target (R1DS1N, Thorlabs). As seen in **Fig 3c** our microscope can clearly resolve two lines separated by 2.2 μm.

**Fig 3.**
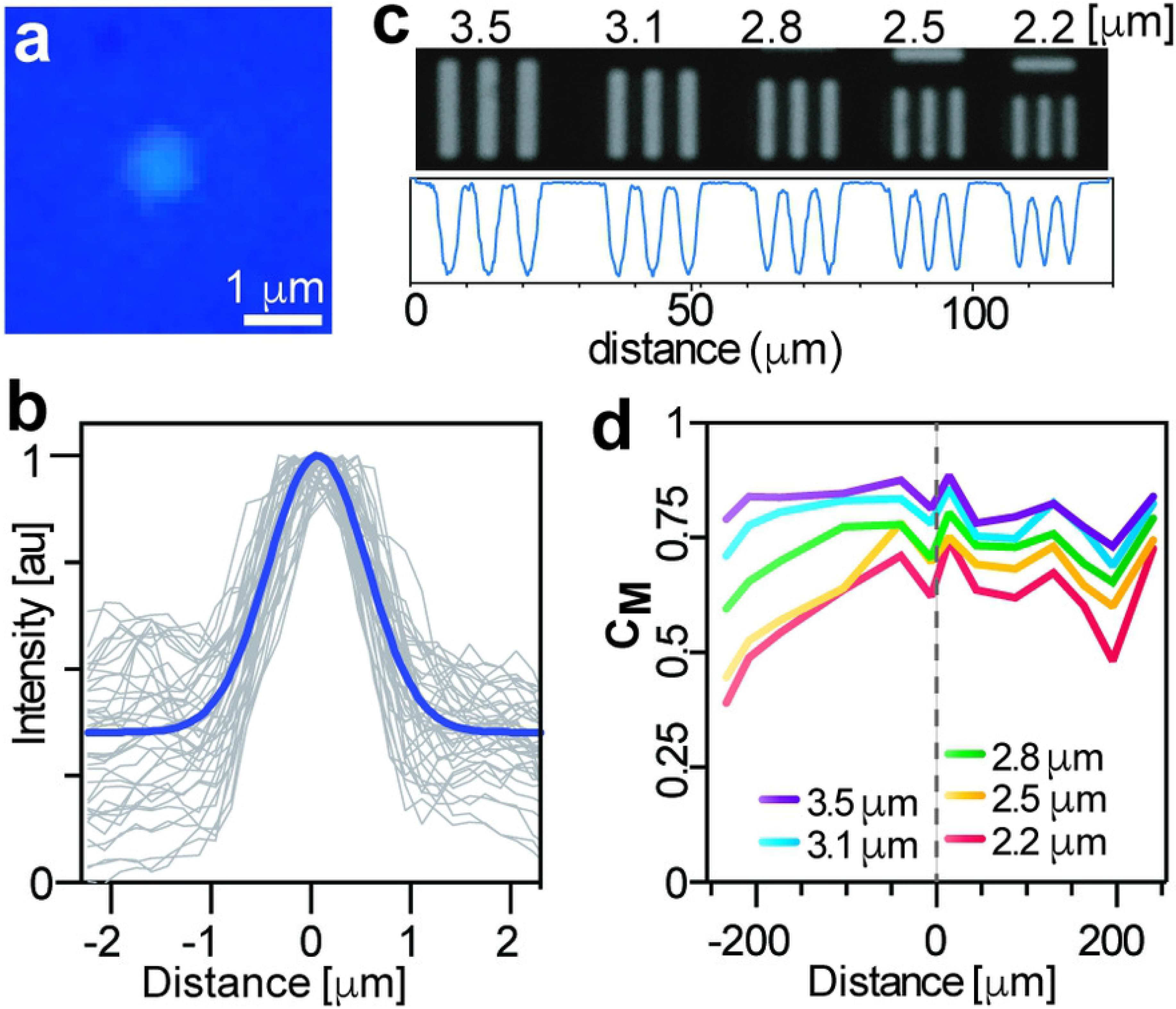
Tests to determine the resolution of the microscope. (**a**) Representative image of a 1 μm fluorescent bead. (**b**) Point spread functions of 40 1-μm beads with the blue line representing the average. (**c**) Brightfield images of the last 5 elements of group 7 on a 1951 USAF resolution target; see full images in S6 Fig. Indicated on top is the thickness of each line. Below each image is the line profile as measured at halfway through all the lines. (**d**) Michelson contrast test using the same USAF target as measured across the width of the image. Gray dashed line represents the center of the image.

A common issue with custom-made microscopes is the radial loss of resolution farther from the center [21]. This is due to the alignment of the optical elements or of the different pieces that comprise the microscope. Using the same USAF target we measured the Michelson contrast, C_M_ (a measurement of contrast [23]), along the width of an image. An element is determined to be resolvable if C_M_ ≥ 0.1 [21]. As shown in **Fig 3d**, the center of the image produces sharper contrasts for all the line sets, though there is a pronounced decreased of C_M_ values on the left side of the image, possibly due to the inclination of the sample stage of the microscope. Nevertheless, all the line sets of the USAF target produce a C_M_ > 0.3 even for patterns as small as 2.2 μm, a sufficient resolution for most cell-based measurements [6,13,16] and for the detection of some parasites [13].

### Microscope Illumination

To correct for uneven illumination intensity from light sources, fluorescence microscopes employ the Koehler illumination, a set of lenses positioned between the light source and the sample[24]. Simpler illumination correction systems to project LED light make use of collimator lenses [2], light-pipes, or elliptical mirrors [25] to generate a parallel beam, or in some cases a diffusor [17] or coupling prisms [2] to guide the beam, thus improving evenness of illumination but not fully correcting it [25]. Poor illumination translates into a poor contrast between the sample fluorescence and the background. However, adding these optical elements adds complexity, cost, and space to any microscopy system.

Thus, we wondered whether direct illumination, where the light source (without any optical elements) is placed near the sample, could give us a uniform illumination. This is a simpler and inefficient configuration as only a small percentage of the light produced by the LEDs reaches the sample [25], but because we used high-radiant power LEDs located very close to the sample we hypothesized that this arrangement would still offer a workable illumination for most applications.

Because the color LEDs could not be placed on top of the sample, as it was already occupied by the white LED, the only positions left where the LEDs could be positioned was in the corners of the lid or at the cardinal points; we decided on the latter as it facilitated the mechanical design. We investigated how uniform was the illumination by illuminating fluorescent solutions (Hoechst, FITC, and Rhodamine) at angles of 0°, 22° and 45° with a 1 s exposure times and employing the plastic filters. (We fabricated three different lids to perform this experiment). **Fig 4** shows the heatmaps of the images captured by the CMOS color sensor at these different angles and considering only the digital channel closer to the emitted light. Data from the contribution of the rest of the RGB components can be found in **S5-S7 Fig**. As it can be observed, vignetting is appreciable in most cases, but it is more manifest at an angle 0°. However, as the angle increases to 22° this vignetting is reduced and almost eliminated for the red (R) channel and to a lesser degree in the green (G) channel, but it is still significant for the blue channel (B): a reduced illumination is noticed on the left side of the heatmap, opposite to where the UV LED was placed. Increasing the angle to 45° reduces this vignetting for the B channel compared to the other angles; however, for the R channel the left side is less illuminated than the right side while the G channel shows a higher radial darkening than at 22°. Emission detection in the other channels is highly attenuated but not completely blocked; this can be attributed to the poor optical quality of the plastic filters. Others [16] have reported that employing the color filter array (CFA) of the CMOS sensor is sufficient to block the excitation light; we did not found this possible perhaps because of the quality of our sensor CFA. However, it is important to highlight that to improve image quality, color and dichroic glass filters can be fitted in our microscope. In general, the most suitable arrangement in our microscope to get the most even illumination is for the UV LED to be located at an angle of 45° and for both the red and blue LEDs placed at an angle of 22°. Note that this characterization would have to be performed if different LEDS are used as they come with different lenses and sizes. In summary, a simple illumination configuration, in which the LEDs are placed in close proximity to the sample, can provide an illumination comparable to critical illumination [25].

**Fig 4.**
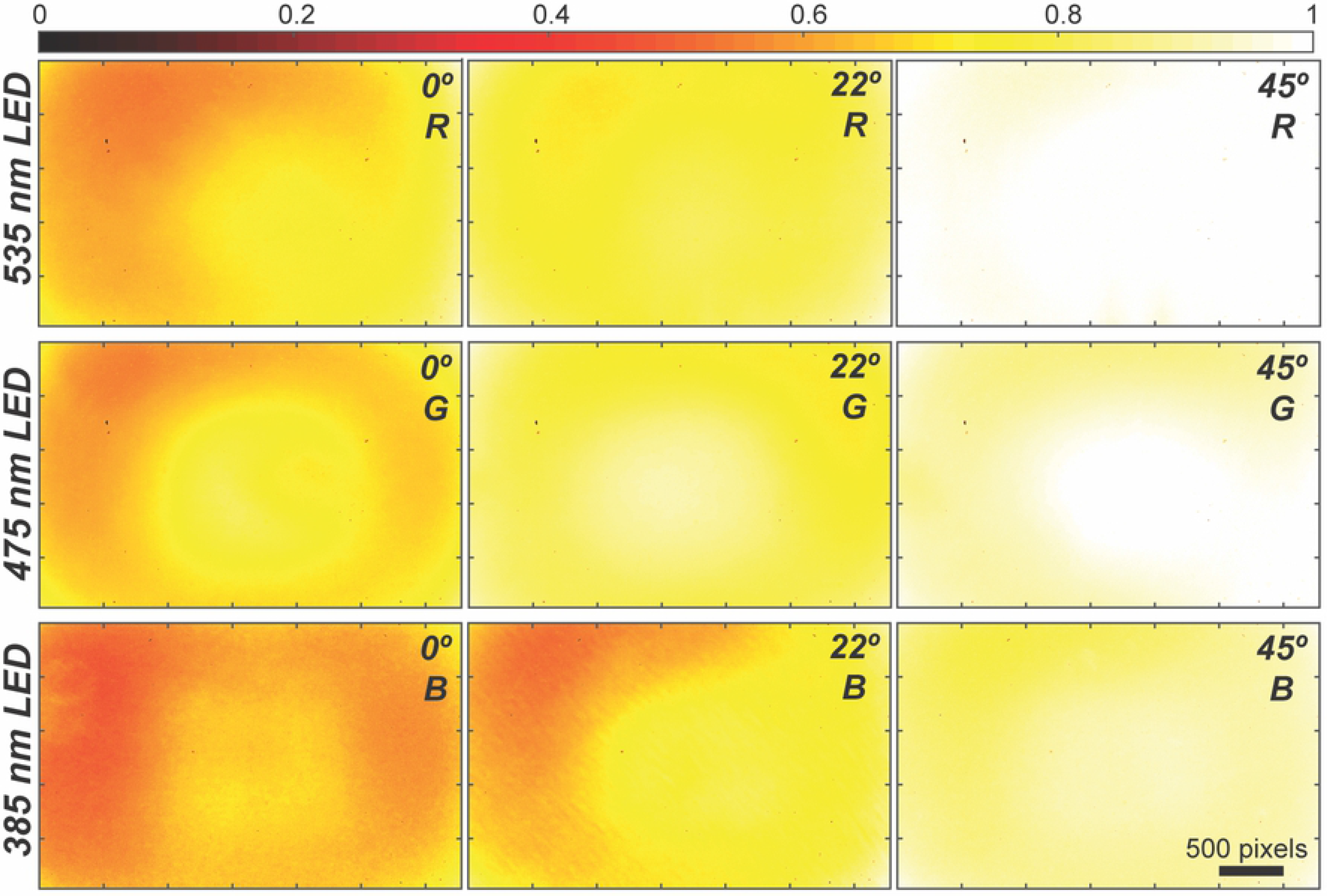
Effects of the angle position of the LEDS on the illumination uniformity using direct illumination. Each row corresponds to a different LED; the angle and the digital color channel analyzed are indicated on the top right corner. R: Red, G: Green, B: Blue. Fluorescence intensity has been normalized to a maximum value of 1.

### Fluorescence images

To demonstrate the utility of our microscope to image single mammalian cells, we acquired bright field and fluorescence images using plastic filters (<$1) and optical glass filters (>$100). **Fig 5a** shows micrographs of THP-1 cells stained with EthD-1 (ex/em 528/617 nm), Calcein-AM (ex/em 496/516 nm) and DAPI (ex/em 360/460 nm) using plastic filters. The intensity profile across the diameter of two cells for the three-fluorescence channels is plotted in **Fig 5b**, after subtracting background intensity. As can be observed, plastic filters block fluorescence bleed-through from neighboring channels. Also, there is no significant difference with commercial glass filters, **S8 Fig**. This experiment demonstrated that our fluorescence microscope produces sufficient image quality to detect and analyze single cells.

**Fig 5.**
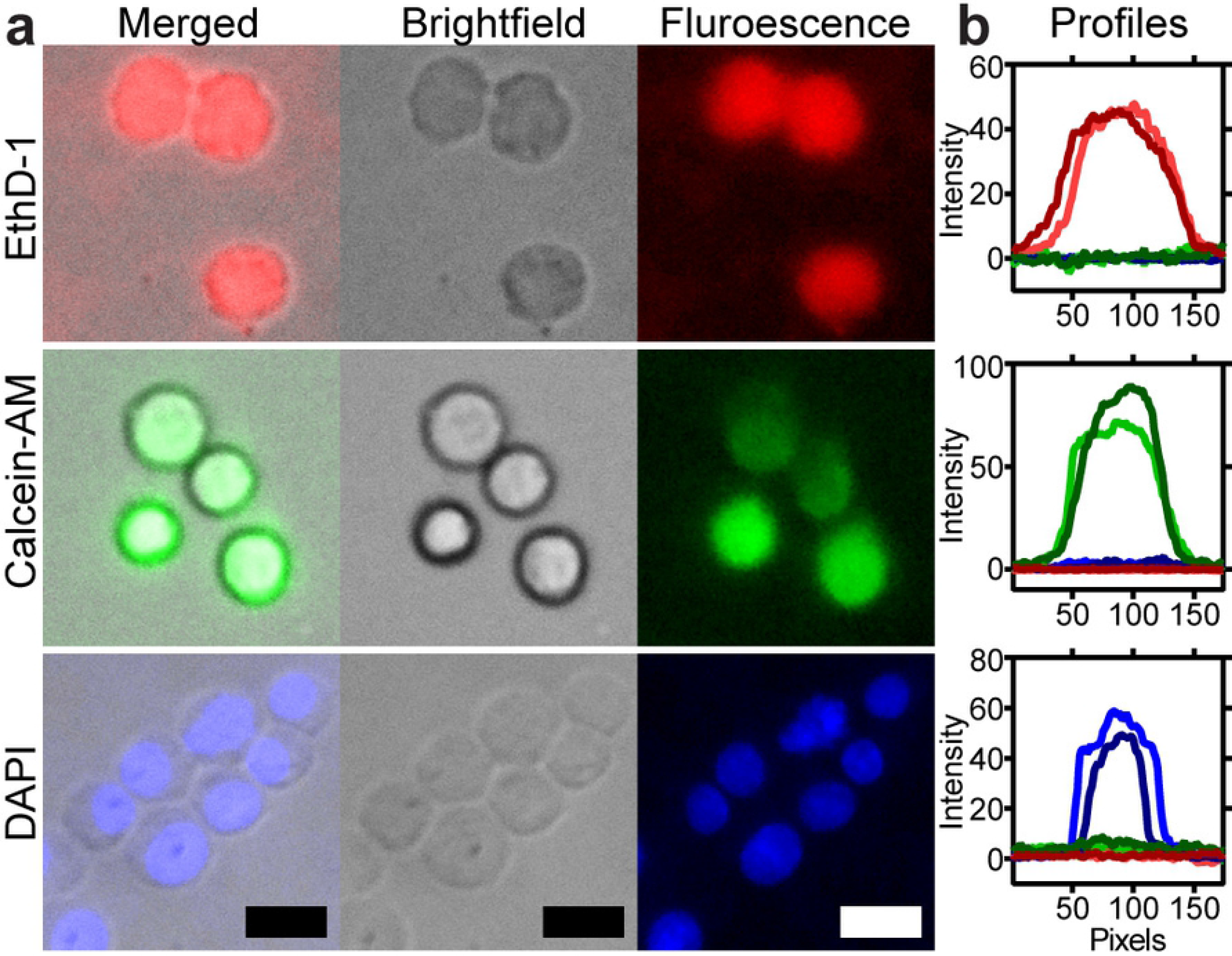
Fluorescence and brightfield micrographs of cells captured with our microscope. (**a**) Brightfield and fluorescence micrographs of THP-1 monocytes stained with three different fluorescent dyes. (**b**) Graphs show the fluorescence intensity profile across two cells in all channels; color represents the fluorescence contribution from each channel. From top to bottom, red, green and blue fluorescence channel. Scale bar = 15 μm.

### Single-cell assay

To demonstrate the utility of our microscope in quantitative biological experiments, we performed a fluorescence time-lapse experiment to track the production of neutrophil extracellular traps (NETs) from single cells [26]. The nuclei of neutrophils isolated from peripheral blood were first stained with Hoechst, and then captured in a PDMS device containing an array of microwells (20 μm in diameter). Next, cells were incubated with both LPS (to stimulate the production of NETs) and with Sytox Orange (a DNA stain impermeant stain).

Brightfield and fluorescence images (UV and Red) were acquired every 10 min for 2 h with our microscope (shown in **Fig 6a**) and with a high-end inverted fluorescence microscope for comparison (not shown). As it can be appreciated it is possible to distinguish single cells trapped in each well. Analysis of individual wells captured with our microscope are shown in **Fig 6b**, each gray trace corresponds to the fluorescence intensity of one individual well while the thick colored curves indicates the average of ∼170 wells. The results for Sytox Orange with both microscopes have similar trends: in both cases most of the wells showed a gradual increase in fluorescence intensity after ∼15 min, reaching a peak at ∼50 min (an indication of loss of plasma membrane integrity and thus yielding an indirect measurement of NET formation), and slowly decreasing as the DNA diffused out of the wells. In the case of Hoechst, the results of our microscope compare favorably to the Zeiss microscope, with both data showing a slight fluorescence intensity from the beginning of the assay —as all the cells nuclei were stained—, increasing gradually to a maximum intensity at 50 min, to eventually decrease afterwards. A negative control experiment where cells were incubated with Hank’s solution did not show any change in fluorescence intensity over time, **S9 Fig**. The slight difference in the data obtained with the Zeiss microscope —higher peaks and smother curves— was expected given the quality of its optical filters and its sophisticated illumination, while ours had a higher background noise. Overall, it is possible to perform quantitative biological experiments in our microscope, enabling fluorescence time-lapse microscopy.

**Fig 6.**
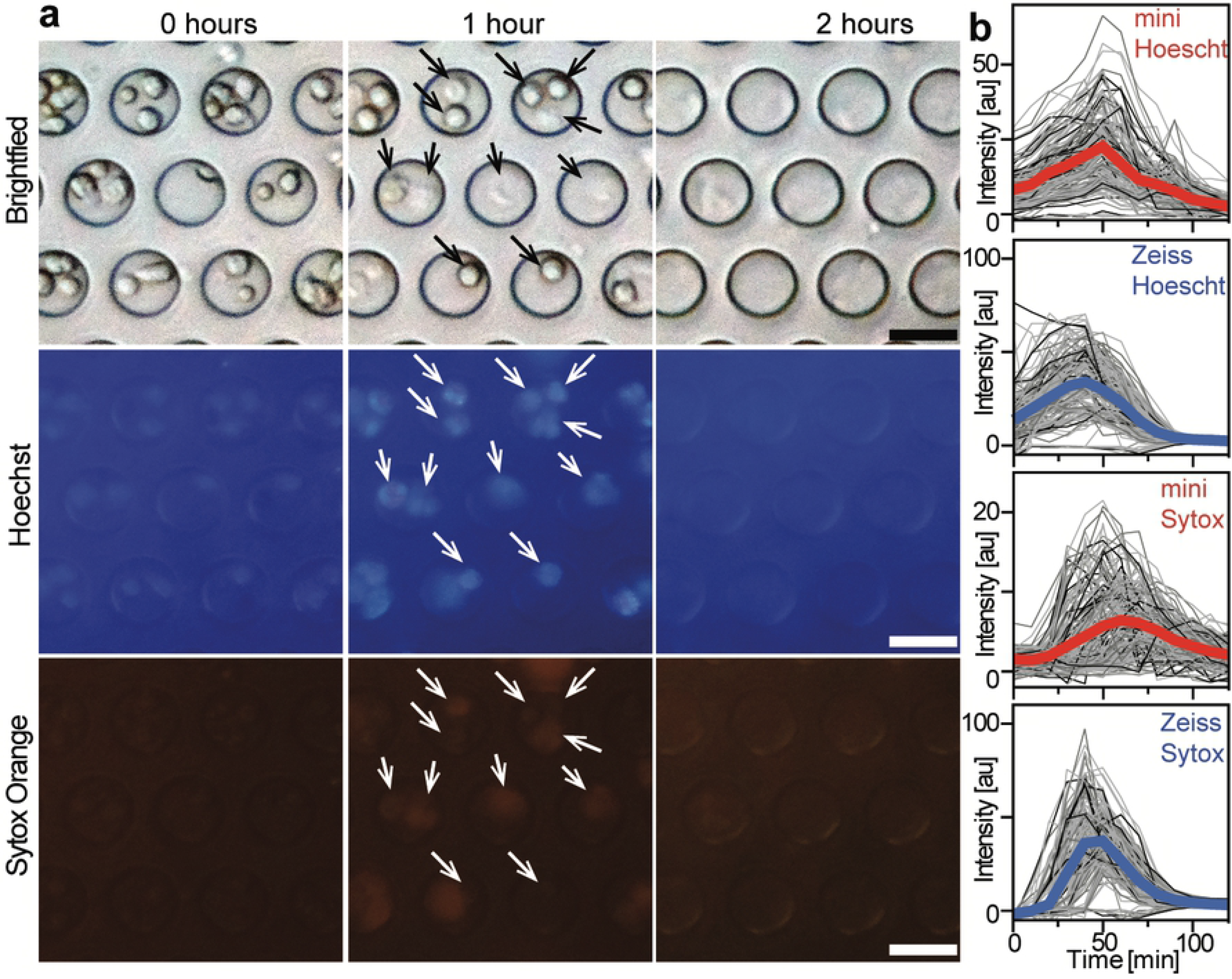
Single cell assays in microwells. (**a**) Representative images of the neutrophil assay at different time points acquired with our microscope using plastic filters. Arrows point to single cells trapped in the microwells. Scale bar: 20 μm. (**b**) Traces of fluorescence intensities from single wells acquired with the miniature microscope (red) and a Zeiss microscope (blue). Thick lines represent the average of ∼170 microwells.

### Cell tracking

Another feature of our miniature microscope is the capability to record video in brightfield and fluorescence mode. Using a single-channel microfluidic device with a width of 40 μm and a height of 20 μm, we injected THP-1 cells stained with Calcein-AM at a speed of ∼150 μm/s, and recorded a video while traveling through the channel (**Figs 7a and 7b**). **Fig 7c** shows a brightfield micrograph of a single cell flowing in the microfluidic channel while **Fig 7d** shows fluorescence micrographs of two cells acquired with a 100 ms exposure time. **S3 Video** shows the facility to focus the optics onto the microfluidic channel and to start recording the cells flowing through the channel, demonstrating the capabilities of our fluorescence microscopy system to not only capture still images of objects sitting on a microscope slide.

**Fig 7.**
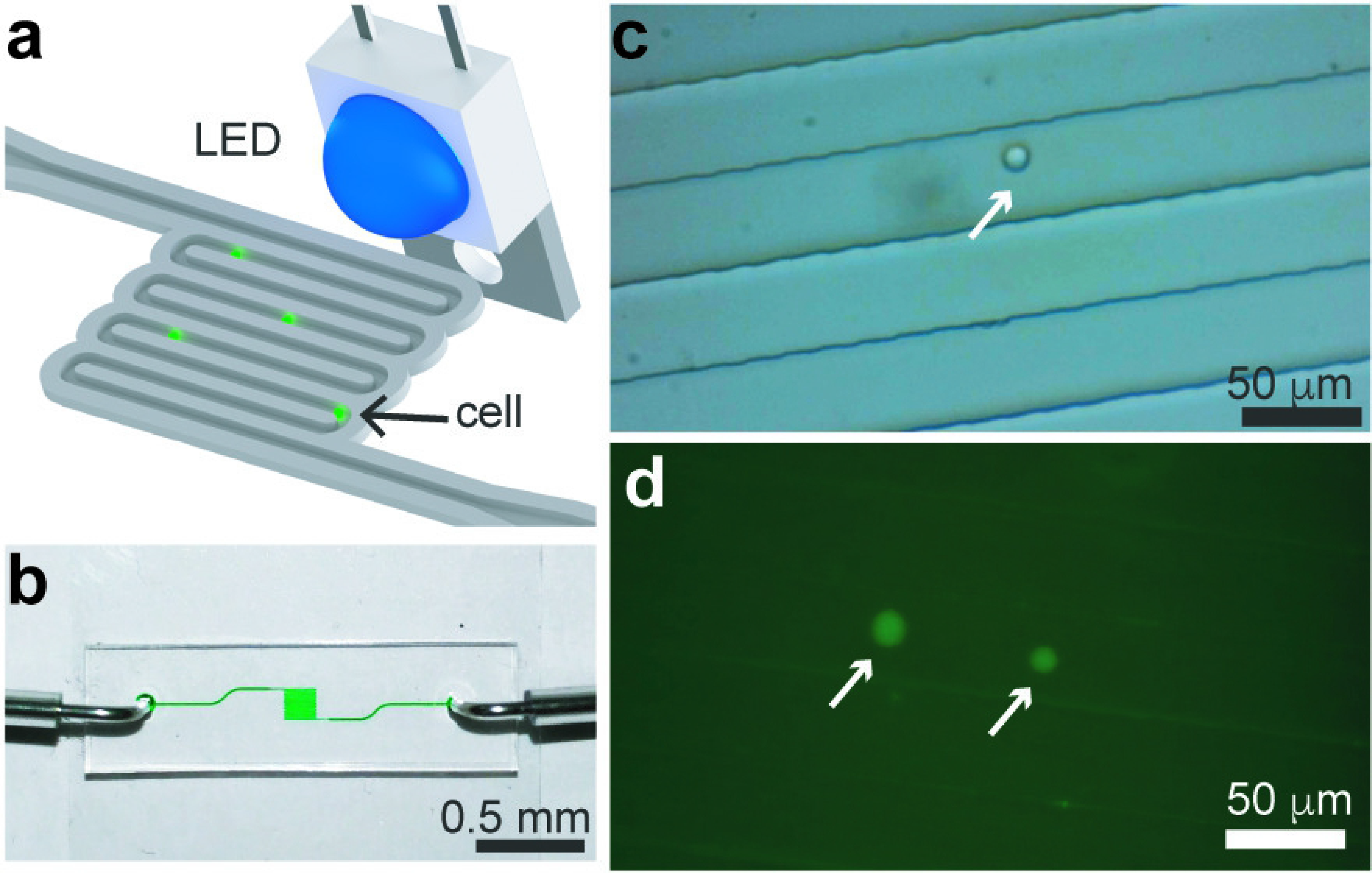
Tracking cells in a microfluidic channel. (**a**) Schematic of the microfluidic device in which cells stained with a fluorescent dye are flowed in a microfluidic device. (**b**) Photograph of the single-channel microfluidic device. (**c**) Brightfield image of a single THP-1 cell flowing at a speed of ∼150 μm/s in a 40-μm wide channel. (**d**) Fluorescence micrographs of two cells flowing inside channel with a 100 ms exposure time. White arrows point to cells.

### Time-lapse microscopy in a cell culture chamber

Because of its size, one of the key features of our microscope is its portability which enables to be used in different settings. To demonstrate this, we used the microscope inside a cell culture incubator set at 37°C with 5% CO_2_ and monitored the culture of THP-1 cells over 26 hours. **S10 Fig** shows a photograph of the miniature microscope placed inside the incubator and the PDMS chamber used for this experiment.

**Fig 8a** shows a series of brightfield micrographs captured every 15 min where it can be appreciated how cells migrate. These images clearly show that cell proliferation is carried out without any disturbance inside the miniature microscope which probes its mechanical robustness. During the time lapse imaging, cells move around the field of view individually or even collectively, staying attached for a few hours after cell division. A sequence of micrographs captured every minute of cells undergoing cell division is shown in **Fig 8b** and **S4 Video**. It is evident that our microscope has enough resolution to observe the lamellipodia of cells extending and retracting, cells migrating over time, and also to distinguish cells undergoing division.

**Fig 8.**
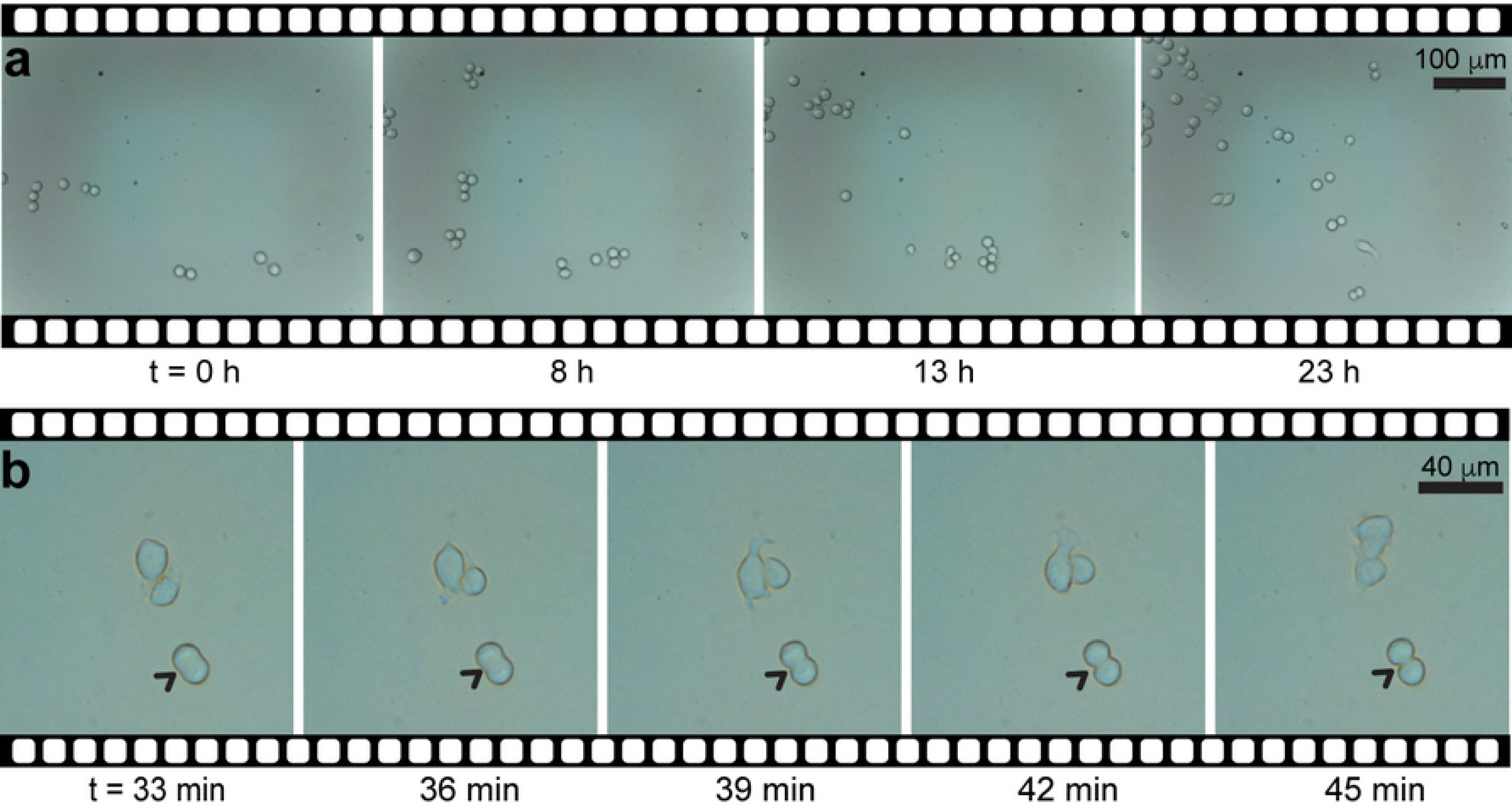
Time-lapse imaging of THP-1 cells. (**a**) Sequence of brightfield micrographs of a cell culture assay that lasted 23 h; images were captured every 15 min. (**b**) Close-up to a sequence of images captured every minute showing the exact moment a cell is dividing (arrows).

## Conclusions

Fluorescence microscopy is an important instrument in biomedical research. Here, we have demonstrated that using 3D printed parts and basic electronic components it is possible to build a miniature 3-channel fluorescence microscope for under $100. In our design, we favored simplicity over other metrics so that it could be assembled rapidly (10 min), although still able to produce sufficient image quality to analyze single cells. We demonstrated that placing color LEDs at different angles from the sample produces a homogenous illumination. We also showed that plastic filters minimized fluorescence bleed-through to the same level of optical glass filters. To demonstrate its application in single cell assays, we monitored the production of NETs from single neutrophils, yielding similar data to a commercial microscope. Our microscope is ideal for downstream applications using microfluidic devices, as demonstrated here, or in situations where space is a premium, for example inside a CO_2_ incubator but we also foresee applications in diagnostics or telemedicine. Because of its low-cost and size, several microscopes could be assembled to monitor several assays at once.

Since LEDs have long-life spans (20,000-50,000 hours) our microscope is suitable for long-term experimentation which could enable acquisition of large amounts of data and translate into better characterization of biological systems. One of the limitations of our microscope are its narrow field of view and its fixed magnification. Thus, an area of opportunity is to develop a motorized stage or use larger CMOS image sensors. Despite this, our microscope performed a fluorescence time-lapse experiment of single cells yielding similar data to a conventional microscope.

## Supporting Information

Electronic Supplementary Information (ESI) available: Detailed fabrication on the microscope parts. Photographs of the step by step assembly of the microscope. Designs of the electronic circuits (PDF). Movies of the microscope assembly and the usage during an experiment showcasing the graphical user interface (MOV).

## Acknowledgments

We would like to thank all the members of the Bio-ARTS Lab at Cinvestav-Monterrey for helpful discussions.

## Supporting information

**S1 Table. Bill of materials of the components used to build the microscope.**

**S1 Video. Assembly of the microscope.**

**S2 Video. Operation of the microscope.**

**S3 Video. Tracking of cells flowing in a microfluidic channel.**

**S4 Video. Time lapse of THP-1 cells growing on a flat surface.**

**S1 Fig. Photograph of the 3D-printed pieces used to assemble the miniature microscope**.

**S2 Fig**. **Design of the microscope tube**. (Left) Front view of the mechanical tube of the microscope. Thread pitch is 1 mm. (Right) Cross-sectional view of the mechanical tube, showing the lens attached to the top of the tube. All dimensions are in mm.

**S3 Fig**. **Schematic and PCB layout of the electronic control unit.** (a) Circuit design. (b) Printed circuit board layout.

**S4 Fig**. **Graphic User Interface (GUI) to control the microscope.** The GUI was coded in Python by combining Tkinter, picamera and OpenCV libraries.

**S5 Fig**. **Analysis of the illumination uniformity at an angle of 0° for all the channels**. Each row corresponds to a different LED; the angle and the digital color channel analyzed are indicated on the top right corner. R: Red, G: Green, B: Blue. Fluorescence intensity has been normalized to a maximum value of 1.

**S6 Fig**. **Analysis of the illumination uniformity at an angle of 22° for all the channels**. Each row corresponds to a different LED; the angle and the digital color channel analyzed are indicated on the top right corner. R: Red, G: Green, B: Blue. Fluorescence intensity has been normalized to a maximum value of 1

**S7 Fig**. **Analysis of the illumination uniformity at an angle of 45° for all the channels**. Each row corresponds to a different LED; the angle and the digital color channel analyzed are indicated on the top right corner. R: Red, G: Green, B: Blue. Fluorescence intensity has been normalized to a maximum value of 1

**S8 Fig**. **Comparison of images captured with both microscopes.** Brightfield and fluorescence micrographs captured with our microscope using plastic filters (left) or Zeiss filters (right). THP-1 cells stained with different fluorochromes: EthD-1 (red), Calcein-AM (green) and DAPI (blue). Images using all filters were acquired for each fluorochrome. Graphs show intensity profile of two cells across all channels, showing that there is no fluorescence bleed-through between channels.

**S9 Fig**. **Negative control experiments.** Traces of fluorescence intensities from single wells for the negative control experiment shown in **Fig 6**.

**S10 Fig**. **Assay inside a cell culture incubator**. (a) Photograph of the inside of a cell culture incubator showing the microscope and the microcomputer Raspberry. (b) Photograph of the cell culture chamber made of PDMS.

